# The evolution of novel biotic interactions at ecological margins in response to climate change involves alleles from across the geographical range of the UK Brown Argus butterfly

**DOI:** 10.1101/2022.02.07.479435

**Authors:** Maaike de Jong, Alexandra Jansen van Rensburg, Samuel Whiteford, Carl J. Yung, Mark Beaumont, Chris Jiggins, Jon Bridle

## Abstract

Understanding the rate and extent to which populations can adapt to novel environments at their ecological margins is fundamental to predicting the persistence of biological communities during ongoing and rapid global change. Recent range expansion in response to climate change in the UK butterfly *Aricia agestis* is associated with the evolution of novel interactions with a larval food plant, and the loss of its ability to use its ancestral larval host species. Using ddRAD analysis of 61210 variable SNPs from 261 females from throughout the UK range of this species, we identify genomic regions at multiple chromosomes that are associated with these evolutionary responses, and their association with demographic history and ecological variation. Gene flow appears widespread throughout the range, despite the apparently fragmented nature of the habitats used by this species. Patterns of haplotype variation between selected and neutral genomic regions suggest that evolution associated with climate adaptation is polygenic, resulting from the independent spread of existing alleles throughout the established range of this species, rather than the colonisation of pre-adapted genotypes from coastal populations. These data suggest that rapid responses to climate change do not depend on the availability of pre-adapted genotypes. Instead, the evolution of novel forms of biotic interaction in *Aricia agestis* has occurred during range expansion, through the assembly of novel genotypes from alleles from multiple localities.

## Introduction

Predicting population and community persistence in the face of a changing and more variable climate remains an urgent priority for biologists (Bridle & van Rensburg 2020). A critical unknown is when and to what extent evolutionary responses will buffer the effects of climate change on ecological communities, or will allow species to shift their ranges to track changes in suitable climate (Hoffmann & Sgrò 2011; Angert *et al*. 2020). Of particular interest is how specialist interactions between species (e.g. parasites and hosts, herbivores and food plants) will limit the availability of suitable habitat, so preventing range shifts (Chen et al. 2011), and how these interactions will change as species encounter novel environmental regimes (Nadeau *et al*. 2017; Hoffmann & Bridle 2021), O’Brien et al. in press).

Population genetic models of adaptation at ecological margins predict that steep and patchy ecological gradients prevent species from tracking changing climate, leading to local (and eventually global) extinction (Polechová & Barton 2015; Bridle *et al*. 2019). Other models of evolution at range margins predict that adaptive shifts are more likely if historic populations have been exposed to variable environments which maintain genetic variation across the species geographical range (Kopp & Matuszewski 2014). In support of this, empirical studies suggest that, although many generalist species have shifted their distributions to track available habitat, almost 75% species with specialist biotic interactions have declined or failed to shift their ranges to match suitable climate (Parmesan 2006; Hill *et al*. 2011).

A key question is whether rapid evolution at ecological margins depends on genotypes already present somewhere in a species’ geographical range. For example, under global climate warming, alleles at the equatorial part of a species’ range may support adaptation at poleward margins, provided gene flow is sufficiently widespread. Such widespread gene flow among marine populations is likely to explain repeated radiations of sticklebacks into freshwater lakes from their marine ancestors (Jones *et al*. 2012). If such pre-existing (“standing”) allelic variation is necessary for rapid evolution, population persistence may depend on maintaining large populations throughout the species’ geographical range, or on translocations from appropriate environments if dispersal among populations is limited (Bridle *et al*. 2009; Hoffmann & Sgrò 2018). By contrast, if rapid evolution depends on novel mutations, then a population’s current size is more relevant in predicting its adaptive potential, rather than its connections with other populations, or its historical size.

Many Lepidoptera species are associated with particular host plant species, particularly at the larval stage, due to plant defence against herbivory, and the particular microclimates that host plants offer as oviposition sites (Jaenike 1990; Stewart *et al*. 2021). Such specialist biotic interactions slow or prevent range expansion into areas that may be climatically suitable, but where the preferred host plant is rare. In the UK, more than 90% of Lepidoptera species that are habitat specialists have contracted their ranges (Warren *et al*. 2001). Limits to habitat availability caused by specialist host plant interactions are also associated with lags in climate responses (Chen *et al*. 2011), with habitat availability explaining 25% of the variation in range expansion rates, even accounting for other factors (Platts *et al*. 2019). Such effects of habitat availability suggest that range shifts in specialists in response to climate change depend on the evolution of novel biotic interactions at ecological margins. Habitat specialists that have tracked changing climates therefore provide an exceptional opportunity to understand the evolutionary responses demanded by ecological gradients made locally steep by particular biotic interactions (Bridle *et al*. 2019; O’Brien et al. <u>in press</u>).

The Brown Argus butterfly, *Aricia agestis* (Polyommatinae: Lycaenidae), is a habitat specialist which has approximately doubled its geographic range in the UK since 1970-1982 (Asher *et al*. 2001) in response to climate change (Thomas *et al*. 2001; Bodsworth 2002; Pateman *et al*. 2012). In mainland Europe, annual plants from the family *Geraniaceae (Geranium* and *Erodium*) are its predominant larval hosts (Tolman 1997; Asher *et al*. 2001). For most of the 20^th^ century in the UK, however *A. agestis* only used *Geraniaceae* as a host plant in a few long-established *A. agestis* populations found in southern and eastern coastal sand dunes (Heath *et al*. 1984). Instead, UK *A. agestis* was largely limited to habitat where the Rockrose host plant (Cistaceae; *Helianthemum nummularium*) is abundant. Rockrose grows only on chalk or limestone soils, and its perennial growth form probably provides a predictable larval microclimate and food supply for *A. agestis*, regardless of annual temperature variation (Stewart et al. in press). By contrast, Geranium annuals are 4-7 times more sparsely distributed across the landscape, and are more affected by seasonal climate, performing well during wet springs and warm summers, but becoming rare and lower quality when springs are dry and summers are wet (Pateman *et al*. 2012; Stewart *et al*. 2021).

These data suggest that for most of the 20^th^ century, *A. agestis* could persist solely in the UK on *Geraniaceae* host plants where south-facing sand dunes provided locally elevated microclimates for rapid larval growth. However, increasing summer temperatures since the 1990s have made non-coastal *Geranium* populations available for sustained occupation by *A. agestis* for the first time, leading to rapid range expansion northward, into areas where Rockrose host plants are rare (Pateman *et al*. 2012; Figure 1).

**Figure 1.**
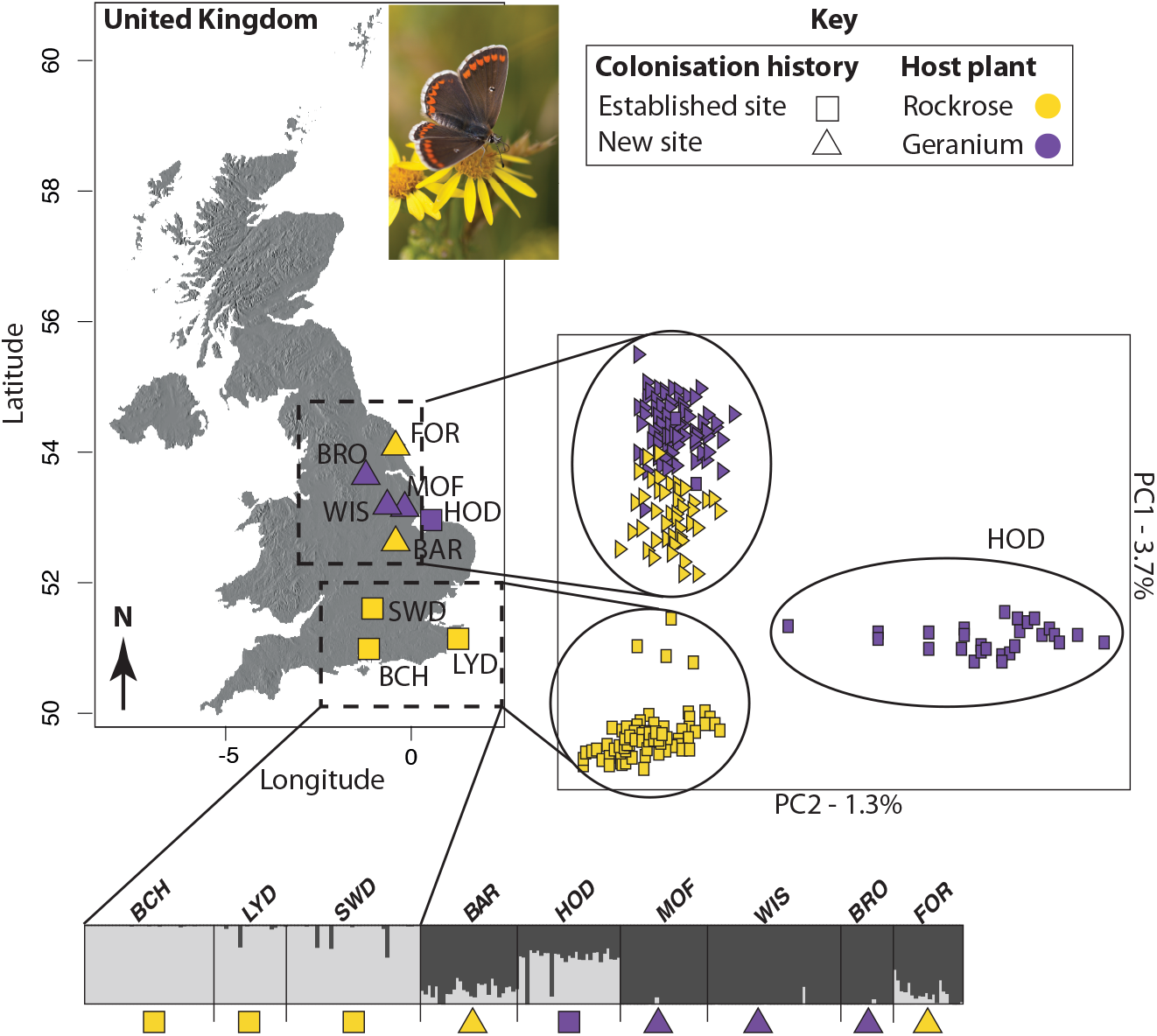
Population structure. Population structure of *A. agestis* sampled across their latitudinal range in the UK. Colonisation history and dominant host plant at each site is shown with shapes and colours. A) The geographic locations of sampled sites. B) Partitioning of genetic variance on a PCA. Note that the graph has been rotated to reflect the latitudinal gradient. PC1 explained 3.6 % of the genetic variance and separates the northern and southern populations. PC2 explained 1.3 % of the variation and separates HOD from the rest of the samples. C) The proportion assignment of each individual to one of two genetic clusters as estimated by fastStructure. Each vertical bar represents an individual, with populations ordered from South to North. Photo credit: Callum Macgregor.

Studies of female oviposition preference and genotyping at AFLP markers demonstrate that range expansion into higher latitude *Geranium* sites has been associated with evolutionary change (Buckley *et al*. 2012). Newly-colonised areas show an increased preference of mothers to oviposit on the widespread *G. molle (Geranium*) (Thomas *et al*. 2001; Bridle et al. 2014), as well as reduced variation in female oviposition preference, increased dispersal ability, and less consistent preference for the locally most common host plant species (Bridle *et al*. 2014). In addition, field transplants of individual females onto natural host plants demonstrate that, although females from Rockrose-dominated (established) sites in the south of the range oviposit on both Rockrose and *Geranium* plants, females from *Geranium*-dominated (newly-colonised) sites north of the range will only oviposit on *Geranium* plants (Buckley & Bridle 2014). This suggests that range expansion driven by climate adaptation in *A. agestis* has been associated with a narrowing of oviposition preference (albeit to exploit a more widely distributed host plant), involving loss of the ability to use its ancestral UK host plants in the newer parts of its range.

The example of the UK Brown Argus suggests that climate-driven range shift by a habitat specialist has required an evolutionary change in species’ interactions. In this case, evolution of host plant has effectively smoothed a steep and patchy ecological gradient at the species’ poleward margin, allowing access to habitats that have recently become climatically suitable, even though they lack the host plants typically used by long-established populations (O’Brien et al. in press). Such a system represents an exceptional opportunity to understand the ecology and genetics of rapid adaptation, and to assess the likelihood and likely context of such adaptation in other species and circumstances, particular for species that depend on particular interactions with other species. In *A. agestis*, a key question is whether its expansion involved the colonisation of novel areas by females from coastal UK populations that already used only *Geraniaceae* food plants. Alternatively, did the observed shift in species’ interactions involve the creation of novel genotypes *in situ*, either from new mutations arising locally, or from selection on standing genetic variation already found in southern UK populations that are able to use both Rockrose and *G. molle* (Buckley and Bridle, 2014).

In this paper, we use genome-wide SNP markers to assess the distribution of genetic variation across the UK, and to test the genetic basis of adaptation at the newly colonised sites. Specifically, we: (1) test for reduced genomic variation associated with range expansion, suggesting selective sweeps at the range margin for specialisation on *Geranium* host plants; 2) identify regions of the genome under selection, and their genomic distribution and likely function; and 3) determine whether evolution during range expansion has occurred through colonisation of existing genotypes that use only *Geranium* host plants, or through the assembly of novel genotypes from alleles sourced from throughout the geographical range.

## Materials and Methods

### Study system and sample collection

Nine *Aricia agestis* populations (276 individuals, 19-45 sampled per site) were sampled in the summers of 2013 and 2014 across their latitudinal distribution in the UK. Populations were chosen to include long established (present since 1970-1982) and newly colonised sites (since 1995-1997; Thomas *et al*. 2001), and sites were classified as either dominated by Geraniaceae (which includes *G. molle, G. dissectum*, and *Erodium cicutarium*) or Cistaceae (Rockrose *Helianthemum nummularium*; Table 1).

**Table 1:**
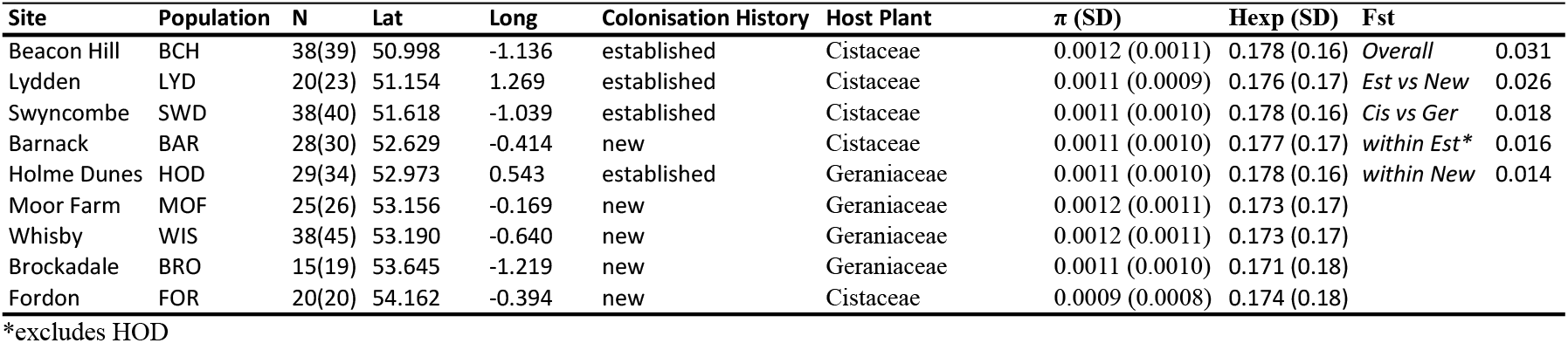
Sample sites and locations. Characteristics of the nine *A. agestis* sites sampled across the UK, ordered from south to north. The number of individuals (N) included in the final analysed dataset is shown, with the number of sequenced individuals in brackets. Sites were defined as established (occupied since 1970-1982) or new (occupied since 1995-1997), and as *Geraniaceae* or *Cistaceae* based on the dominant host plant at that site. We report two measures of genetic diversity; nucleotide diversity (π) and expected heterozygosity (Hexp; see Figure S1) as the mean and standard deviation (SD) estimated across all neutral SNPs. Nei’s F_ST_ was estimated for between all nine populations (*Overall*), and between the Colonisation History (*Est vs New*) and Host Plant (*Cis vs Ger*) groups, as well as between populations within the Established (*within Est*) and New sites (*within New*). HOD was excluded from the *within Est* comparison given its geographic distance from the established South.

### Generating genome-wide markers for population genetics

DNA was extracted from the head and half of the thorax each of 276 individuals using a Qiagen DNeasy Blood and Tissue kit. DNA was eluted in EB buffer and quantified using Qubit 2.0 fluorometer with the DNA BR assay kit (Life Technologies).

Genome wide markers were generated using a modified double digest restriction associated DNA (ddRAD) protocol (Peterson *et al*. 2012). Briefly, genomic DNA of each individual was digested with *PstI* and *EcoRI* restriction enzymes. In total 276 individuals from nine populations were sequenced across six ddRAD libraries. Individuals from a single population were sequenced in at least four different libraries to avoid confounding population structure with differences in library preparation and sequencing between libraries. Each library comprised 48 individuals uniquely identified using a 6-bp DNA barcode. Libraries were sequenced on an Illumina HiSeq 2000 instrument to generate 150-bp paired-end sequences from insert sizes of 300-450 bp.

### Bioinformatic analysis

Raw data were demultiplexed based on individual barcodes using ipyRAD v. 0.7.28 (Eaton 2014) and adapters were removed using Trimmomatic v. 0.36 (Bolger *et al*. 2014). A long-read based reference genome has been assembled by the Sanger Institute and is available under accession number GCA_905147365.1 from the National Center for Biotechnology Information (NCBI; www.ncbi.nlm.nih.gov). To call variants we first mapped the demultiplexed data to the reference genome using BWA-mem (Li 2013). Next, we called variants using mapped read pairs with a PHRED scaled mapping quality higher than 20. We used the SAMtools v.1.8 (Li *et al*. 2009) *mpileup* and BCFtools v. 1.8 (Li 2011) *call* functions to call variants simultaneously across all individuals. Raw individual variant calls were output as a VCF and filtered before downstream analyses using VCFtools v.0.017 (Danecek *et al*. 2011), BCFtools v. 1.8, and PLINK 2.0 (Purcell *et al*. 2007).

The following filters were applied: 1) We retained only variants called with a genotype likelihood (QUAL) PHRED score of more than 20. 2) A minimum mean depth filter of 6x was applied to reduce spurious heterozygote calls. 3) A maximum mean depth of the mean plus twice the standard deviation (646x) was applied to remove duplicate loci. 4) Kinship coefficients were estimated between individuals within each population using the KING method (Manichaikul *et al*. 2010). Individuals were removed to exclude any second order or higher relatives (phi > 0.05). 5) Individuals with a genotyping rate of less than 60% were removed. 6) To include loci that were evenly genotyped across all populations, we excluded loci genotyped in less than 50% of individuals within each population. 7) We allowed a global minimum minor allele frequency (MAF) of 1%. The final dataset comprised 251 individuals genotyped at 61210 variants.

#### (1) Changes in genomic variation associated with range expansion

##### Population structure and genetic variation associated with range expansion

We assessed population structure and genetic diversity across the sampled range using three complementary methods. First, we visually examined the extent of genetic differentiation between populations using a Principal Component Analysis (PCA) using the R package PCAdapt (Luu *et al*. 2016). Secondly, we estimated the most likely number of genetic clusters with fastStructure (Raj *et al*. 2014). fastStructure uses variational Bayesian inference to approximate the log-marginal likelihood of the data. This approach is attractive for large genomic datasets because it is very rapid compared to a traditional Bayesian approach implemented in Structure (Pritchard *et al*. 2000). We estimated ancestry proportions for up to 10 clusters (K=2-10) using a simple prior on the model. Thirdly, we estimated individual co-ancestry based on a pairwise comparison between samples using a Markov chain Monte Carlo (MCMC) coalescence model implemented in fineRADstructure (Malinsky *et al*. 2018). fineRADstructure estimates coancestry between individuals by comparing haplotypes across all individuals to estimate the nearest neighbour for each locus. Coancestry is divided equally between all individuals with the same haplotype, or between individuals that are the nearest neighbour of a rare haplotype. In this way rare haplotypes and their nearest neighbours receive a higher coancestry weighting than more prevalent haplotypes. Given that rare mutations are on average expected to be of more recent origin than haplotypes that occur at higher frequencies in the population, fineRADstructure is able to estimate recent coancestry in the dataset. No population prior was specified for the analysis. We ran the analysis with a burn-in of 100’000 iterations, followed by 100’000 MCMC steps. RADpainter (Malinsky *et al*. 2018) was used to infer the coancestry matrix and assign individuals to populations using default parameters.

To determine whether genetic divergence between populations increases on average with geographic distance, we estimated population pairwise F_ST_ (Nei 1973) from a subset of unlinked loci using the R package *adegenet* (Jombart *et al*. 2008). We estimated Pearson’s correlation coefficient between genetic distance and log transformed geographic distance, difference in dominant host plant, and difference in colonisation history. Significance was tested with a Mantel test in the *vegan* package (Oksanen *et al*. 2015) in R.

##### Testing for changes in host plant use at the range edge

We tested for genome-wide signals of specialisation on the most prevalent host plant at each site by determining how much genetic variance can be explained by 1) host plant prevalence and 2) site colonisation history. The initial model (Basic Model) determined how much of the genetic variance could be explained by all the variables combined using a redundancy analysis (RDA) as implemented in the vegan package (Oksanen *et al*. 2015) in R. The full Basic Model was GeneticData [MAF matrix] ~ Latitude + Longitude + Host Plant + Colonisation History. Next, we determined the best model to explain variance in the genetic data by removing one non-significant variable at a time based on an automatic permutation of the model and selecting the best variables in each case based on Akaike’s Information Criterion (AIC). This was implemented using the *ordistep* function from *vegan* in R. We then used two partial RDA analyses to estimate the variance explained by Host Plant or Colonisation History when Latitude and Longitude are kept constant.

Previous data on UK *Aricia agestis* suggests that populations expanding at the range margin lost genetic diversity due to the spread of genotypes laying only on Geraniaceae hosts rather than on Geraniaceae and rockrose hosts (Bridle *et al*. 2014; Buckley & Bridle 2014). We tested for a signal of a genome-wide reduction in genetic variation associated with range expansion by comparing gene diversity and mean genome-wide nucleotide diversity between sites dominated by the different host plants or with different colonisation histories. Nucleotide diversity was calculated across 1kb windows within each population using VCFtools v.0.1.14. Expected heterozygosity was calculated using the *basic.stats* function in the R package *hierfstat* (Goudet 2005). We tested whether these estimates differed significantly between established and new populations, as well as between Geraniaceae and Cistaceae dominated sites using a Kruskal-Wallis rank sum test implemented in R. We tested the robustness of these pairwise comparisons using two methods: 1) a jackknife approach, where we repeated the analysis while removing each population in turn, and 2) we randomised the colonisation history and host plant variables and repeated the Kruskal-Wallis rank sum tests.

#### (2) Identifying genomic regions under selection and their geographical distribution

##### Identifying genomic regions under selection across the UK

We used the F_ST_ outlier approach implemented in BayeScan v2.1 (Foll & Gaggiotti 2008) to identify adaptive genotypes associated with 1) different host plants and 2) site colonisation history. This model decomposes F_ST_ between populations into a population (beta) and locus (alpha) component. A locus is deemed to be under selection if the alpha component is needed to explain the diversity at that locus. BayeScan implements a reverse-jump MCMC to explore the parameter space with and without the alpha parameter included, and therefore estimates a posterior probability associated with each model. To identify and exclude any loci associated with a single population, we repeated the analysis with each population removed in turn. Loci that were identified with a false discovery rate of 0.05 in the full analysis and at least two thirds of the jackknife analyses were included in the final set of outlier loci.

##### Identifying the function of outliers

We annotated the outlier loci with snpEff (Cingolani *et al*. 2012) using a database built from the reference genome and annotation available on NCBI (GCA_905147365.1). We used the “closest” function to identify the closest gene to each outlier locus. The gene name and biological process was obtained by searching the UniProtKB database (The UniProt Consortium 2021) for the protein identified in each case.

#### (3) Determining the geographical and colonisation history of evolutionary responses

##### Haplotype networks of outlier loci associated with host plant variation

To determine if alleles associated with adaptation to different host plants spread independently, or as genotypes during population expansion, we constructed haplotype networks for each of the loci by identifying and phasing all outlier ddRAD tags. We subset the mapped reads (bam files) by extracting the outlier variants flanked by 300bp on either side to include all variants within the RAD tag. We kept only sequences with at least two variant sites. We phased each locus with WhatsHap (Patterson *et al*. 2015), which uses sequence data from each individual’s bam files to inform phasing. All unphased sites were removed. We extracted the haplotype sequences in fasta format and constructed haplotype networks for each locus using the Pegas package (Paradis 2010) in R.

To compare the spread of adaptive versus neutral loci across the landscape we used the same approach to extract a random set of 20 neutral sequences. After removing sequences with less than two variant sites we reconstructed haplotype networks for the 11 remaining sequences.

##### Reconstructing colonisation history using a coalescent approach

The evolution of *A. agestis* to use only *Geraniaceae* at newly-colonised sites could have occurred through the arrival of pre-adapted genotypes that already specialised on these host plants, or through evolution *in situ* at the range edge. The established north-eastern population (HOD) is dominated by *Geraniaceae* and its geographic proximity to newly-colonised sites makes it a potential local source for such pre-adapted genotypes. To test this idea, and to determine the most likely origin of colonists at the new sites, we constructed six demographic models scenarios using the coalescent simulator fastSimCoal2 (Excoffier *et al*. 2013). We compared three models of demographic history with two different migration scenarios applied to each: Model 1) the established southern populations (South) as the source; Model 2) HOD as the source; Model 3) Secondary contact between HOD and South followed by the colonisation of the new sites (Figure 2). We applied two migration scenarios to each model for a total of six models: a) a full migration matrix of asymmetric gene flow, and b) a complete absence of gene flow after divergence. We removed all non-neutral loci identified by our tests for selection and excluded the Z-chromosome from our analysis.

**Figure 2.**
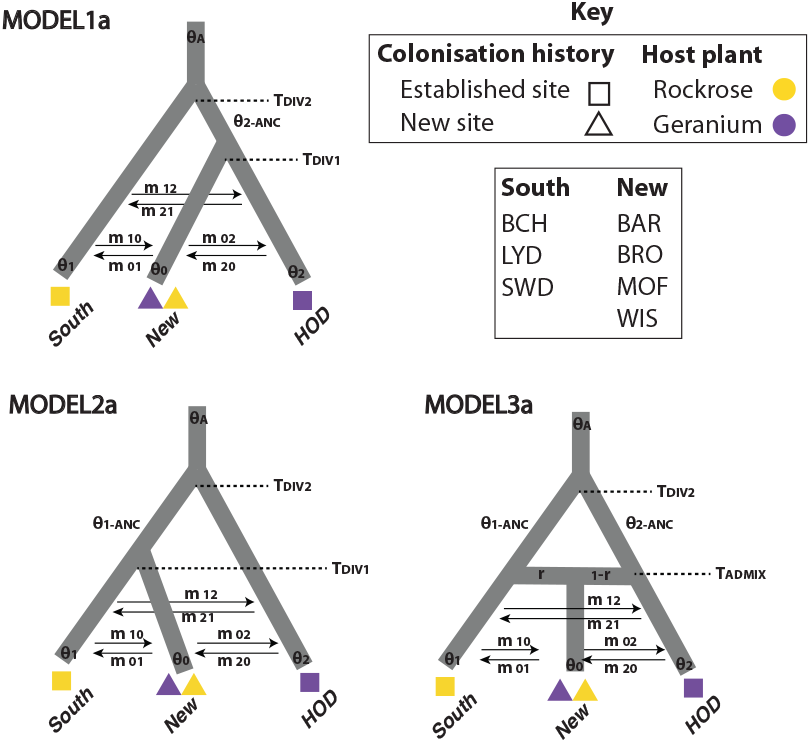
Demographic models tested with fastSimCoal2. Three demographic models were compared using FastSimCoal2: Model 1) Colonisation of the new sites exclusively from HOD followed by gene flow between all populations, Model 2) Colonisation of new sites exclusively from the South followed by gene flow between all populations, Model 3) Secondary contact and subsequent colonisation of the new sites by admixed populations from HOD and the South. All models were tested with two different migration matrices: a) assymetric gene-flow after divergence, b) no gene flow. Parameters included in the model were mutation-scaled effective population size (θ), migration rates per generation (*m*), the time of divergence (T_DIV_) or admixture (T_ADM_) between populations, and the proportion of the source population transferred to the sink population (*r*).

Missingness is a characteristic of reduced representation libraries because loci sequenced in different libraries don’t overlap exactly. However, if we minimise the missingness in the dataset, too few loci remain to accurately estimate the site frequency spectrum. At the same time, including sites with a high proportion of missingness can bias the estimated site frequency spectrum. Therefore, to increase the number of loci in the final dataset, we used resampled data within populations to reduce missingness. This approach resamples loci that have a user-specified minimum number of genotypes across all samples. In this way, missingness within an individual is circumvented and a full genotype matrix can be created. Our final dataset comprised 9735 SNPs. Scripts to downsample the data and construct the minor allele frequency spectrum were obtained from Vitor Sousa, University of Lisbon (https://github.com/vsousa/EG_cE3c/tree/master/CustomScripts/Fastsimcoal_VCFtoSFS).

The effective population size was fixed for “South” in the model so that all parameters could be estimated relative to this value. The effective population size (*N_e_*) was calculated from the mutation rate (μ) and nucleotide diversity (π). Nucleotide diversity was calculated across all variant and invariant sites in windows of 1kb using vcftools (--window-pi function; π=0.001). We assume a mutation rate of 2.9 × 10^−9^ per base per haploid genome per generation based on the only direct estimate of Lepidoptera mutation rates (*H. melpomone*, Keightley *et al*. 2014). The median effective population size was calculated as *N_e_* = (π/4μ); South=86207 (range 898-712069). For parameter estimation we assume a generation time of 0.5 years, because *A. agestis* in Britain are bivoltine.

Given the uncertainty associated with these parameters, and the small site frequency spectrum compared with the number of parameters estimated (9-15), we expect wide and overlapping confidence intervals and uncertainty in the parameter estimates. This meant that we used the test to determine the relative likelihood of the different demographic models, rather than the absolute values of the estimated parameters. We ran 100 independent simulations of each model in fastSimCoal2. Each run comprised 100 000 coalescent simulations and 40 expectation maximisation cycles. All parameters and priors are documented in Table S4. We evaluated the model fit based on the lowest log likelihood and Akaike’s information criterion (AIC). To compare the models directly we rescaled AIC as the difference between the lowest AIC and the AIC for each competing models. This is shown in the results as ΔAIC, where the best model has a ΔAIC of 0 (Table 4). The point estimates of each demographic parameter for the best supported model were obtained from the run with the highest composite maximum likelihood score.

Confidence intervals (CI) were estimated for the parameters by simulating 100 site frequency spectra using the maximum likelihood point estimates for the best run. Parameter estimates were re-estimated using 100 independent simulations of the model for each of the simulated site frequency spectra The lower and upper CI bounds were obtained from the lowest and highest composite maximum likelihood estimates obtained from the 100 iterations.

## Results

After applying filters, the final datasets comprised 251 individuals from nine populations (n=15-38; Table1) genotyped at 61210 variants.

### (1) Changes in genomic variation associated with range expansion

#### Population structure is associated with latitude

Our results suggest that *A. agestis* populations are largely structured latitudinally, following the likely colonisation route northwards, with the exception of FOR (see below). Overall gene flow was high (F_ST_=0.031 between all sites; Table 1, Table S1), but genetic distance increased significantly with geographic distance (Mantel’s r = 0.78, *p* = 0.001). Similarly, genetic divergence was low (F_ST_=0.026) but significant between established and new sites (Mantel’s r = 0.45, *p* = 0.04), with estimates similar to previous results based on 409 AFLP markers (F_ST_=0.025; Buckley *et al*. 2012). By contrast, genetic divergence was not significant between sites dominated by different host plants (F_ST_=0.018; Mantel’s r = 0.29, *p* = 0.07).

Our PCA analysis showed that the most genetic variance was explained by the divergence between northern and southern populations (PCA1 = 3.7%; Figure 1), with FOR more closely related to the southern populations than the other northern populations, and HOD more differentiated than the others. Similarly, fastStructure results supported two genetic clusters (K=2) that also correspond to the northern and southern populations (Figure 1). Both analyses showed that HOD (the only established population where *Geraniaceae* is the most prevalent host plant) is differentiated from the rest of the UK sites. In the PCA, the divergence between HOD and the rest of the populations explains 1.3% of the total genomic variance (PC2). Results from fastStructure also reveal that HOD and BAR (a newly established site) are genetically intermediate to the northern and southern clusters (Figure 1).

Recent coancestry as estimated with fineRADstructure revealed a more complex relationship between populations. Two well-supported groups were recovered that correspond to the northern and southern populations. However, HOD clustered with the three established southern populations (Figure 3). Substructure within the northern cluster supported differentiation between the newly established Geraniaceae dominated sites and the two sites where Cistaceae are the most prevalent. The Cistaceae dominated sites (FOR and BAR) are geographically well separated, and the Geraniaceae sites (BRO, WIS, MOF) are found between them. This suggests that the new sites were colonised from the established Cistaceae sites, rather than the established Geraniaceae site to the east (HOD). There is also evidence of colonisation through infilling from neighbouring populations, with at least five individuals in the new Geraniaceae sites resembling the neighbouring new Cistaceae and established Geraniaceae haplotypes (Figure 3).

**Figure 3.**
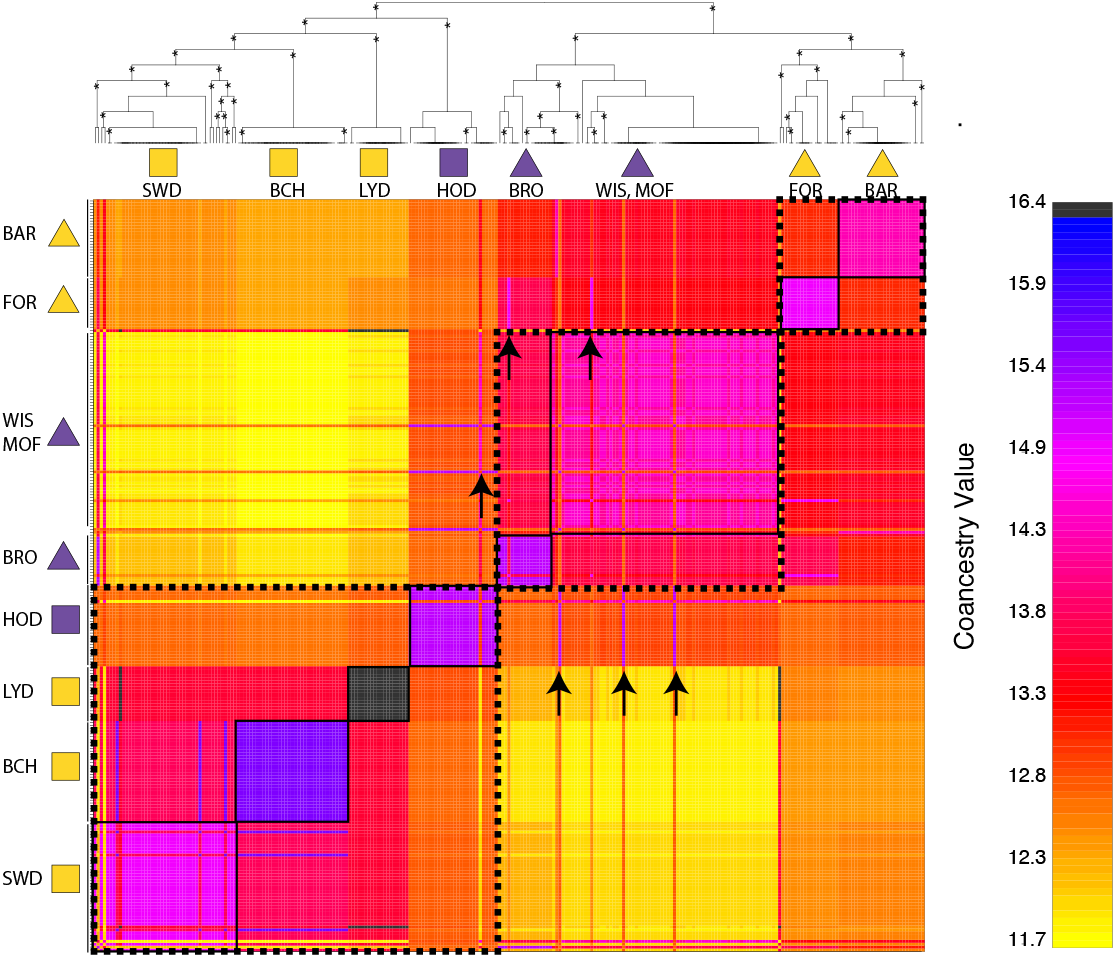
fineRADstructure. Individual coancestry matrix estimated with fineRADstructure and clustered by population using RADpainter. The level of co-ancestry is indicated with colour (high=black/blue; low=yellow) as shown by the colour bar to the right. The maximum a posteriori (MAP) tree shows the inferred ancestry of each individual. Posterior probability branch support above 0.85 is indicated with an asterisk. The inferred co-ancestry groups largely correspond to geographic populations, although there is evidence of haplotypes moving from the new Rockrose sites (BAR and FOR) to the new Geranium sites (WIS, MOF and BRO), as well between the new Geranium sites and HOD. These are indicated with arrows. Hierarchical structure in the data is shown with solid and dashed lines around grouped populations.

#### Genomic variation is slightly reduced in newly-colonised regions

Our results show that there is genome-wide differentiation between new and established sites, and between Geraniaceae and Cistaceae dominated sites, but that the variance in the data cannot be significantly explained by either of these variables, independent of spatial variables (Table 2). The Basic Model explained a large and significant proportion of variance in the genomic data (Basic Model: 66%, p=0.001). The best model retained Latitude and Colonisation History as significant variables. Using partial redundancy analyses we found a large proportion of constrained variance explained by HostPlant use (10%) and Colonisation History (14%), but neither were significant in the Basic Model.

**Table 2.**
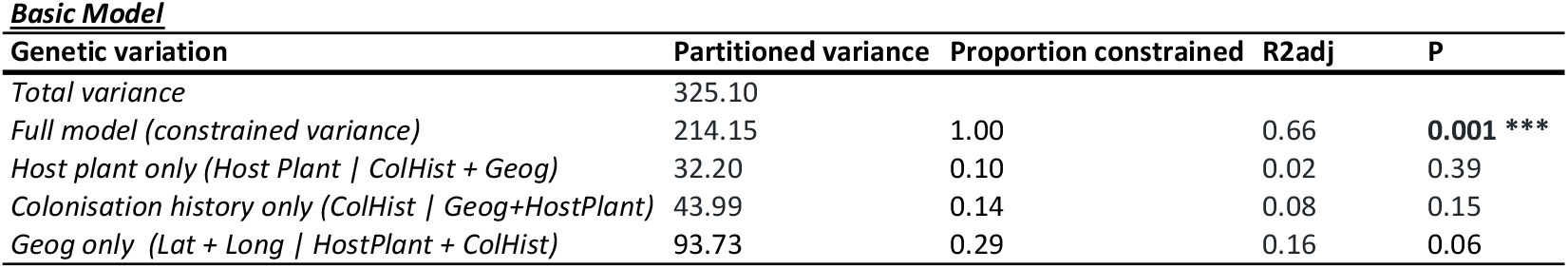
Redundancy Analysis. Partitioning of genetic variance in each geographic transect using RDA and partial RDA analyses. The column “Partitioned variance” shows the total variance of the genetic data (Total variance), the proportion of variance that could be explained by the full RDA model which includes HostPlant, Colonisation History, Longitude and Latitude as explanatory variables (Basic Model), and the proportion of total variance explained by the partial RDA in each case. The column “Proportion constrained” shows the variation explained by each model relative to the total explainable variance. The fit (R^2^adj) and significance (P) of each model is shown, and with significant p-values shown in bold.

Genetic diversity was marginally but significantly lower in newly colonised sites when compared with established sites (Table 1, Figure S1; median Hs, π: new = 0.12, 0.00074; established = 0.13, 0.00077), and in Geraniaceae dominated sites compared with Cistaceae sites (median Hs, π: Geraniaceae = 0.12, 0.00074; Cistaceae = 0.13, 0.00077). For comparisons based on nucleotide diversity the test remained highly significant (*p* < 0.001) except with the removal of MOF or FOR. Randomisation of the host plant or colonisation history variable resulted in non-significant differences. Allelic diversity was significantly different in all cases (*p* < 2.2e-16), but was still significant when the variables were randomised (*p* = 0.026).

### (2) Identifying genomic regions under selection and their distribution

#### Identification and putative origin of loci under selection

We identified 12 loci (137 SNPs; 0.22% of total variants) associated with host plant preference, and 19 loci (239 SNPs; 0.39%) associated with colonisation history. Two loci each on the Z-chromosome and Chromosome 9 were identified as outliers in both “colonization history” and “host plant use” datasets (Table S2). Of the 25 candidate loci that could be assigned an annotation, 21 were located in the intron of a gene, and the remaining four were in intergenic regions 1287-71461 bp from the closest gene. The loci were distributed across the genome, occurring on 8 (Host Plant) and 11 chromosomes (Colonisation History) (Figure 4). The outlier locus with the highest difference in allele frequencies in both cases is a locus on chromosome 9 with pairwise F_ST_ > 0.4. This locus was identified as being under strong selection both in the Host Plant choice and Colonisation History comparison.

**Figure 4.**
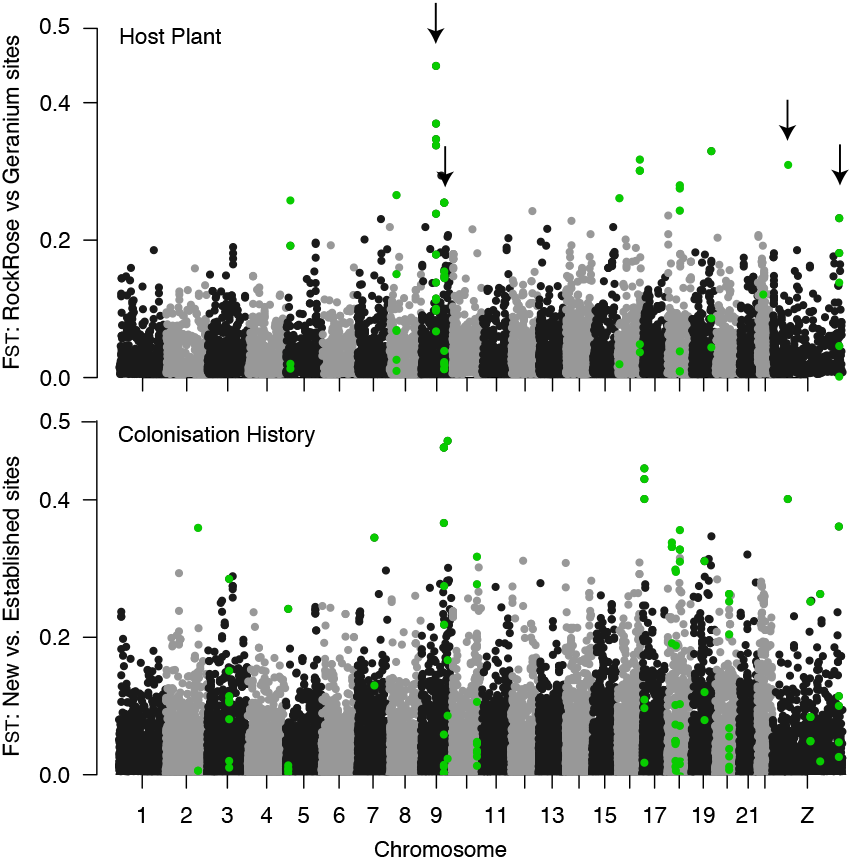
Manhattan plot. The per locus distribution of F_ST_ across the genome between populations defined by A) Host Plant preference, and B) Colonisation history. Outlier loci are coloured green. Four outlier loci, indicated by arrows, were associated with both host plant choice and colonisation history.

### (3) Determining the geographical and colonisation history of evolutionary responses

#### Haplotype networks of outlier loci associated with host plant variation

Haplotype networks of the HostPlant outlier loci (Figure 5; Figure S2) show that haplotypes did not cluster by phenotype, as would be expected where variation in traits is determined by only a few loci (Van Belleghem *et al*. 2018; Kautt *et al*. 2020). This suggests that selection on variation in Host Plant preference acts across many loci with high levels of recombination among them. We also found no evidence that adaptation to Geraniaceae occurred through the rapid influx of pre-adapted alleles (or haplotypes) from HOD, given that no selected haplotype currently found in new Geraniaceae sites was derived from the established coastal Geraniaceae site (HOD). Instead, we found that the Geraniaceae-preferring haplotypes in newly-colonised areas were similar to common haplotypes that occur in both the South and HOD, or only in the South (Table S3). In addition, haplotype networks of neutral loci were indistinguishable from the adaptive loci, which suggests that the adaptive and neutral alleles spread to the new Geraniaceae populations from both the established populations in the South (that can use both *Geranium* and rockrose) and from coastal populations using Geraniaceae, across sufficient numbers of generations for recombination to occur between them.

**Figure 5.**
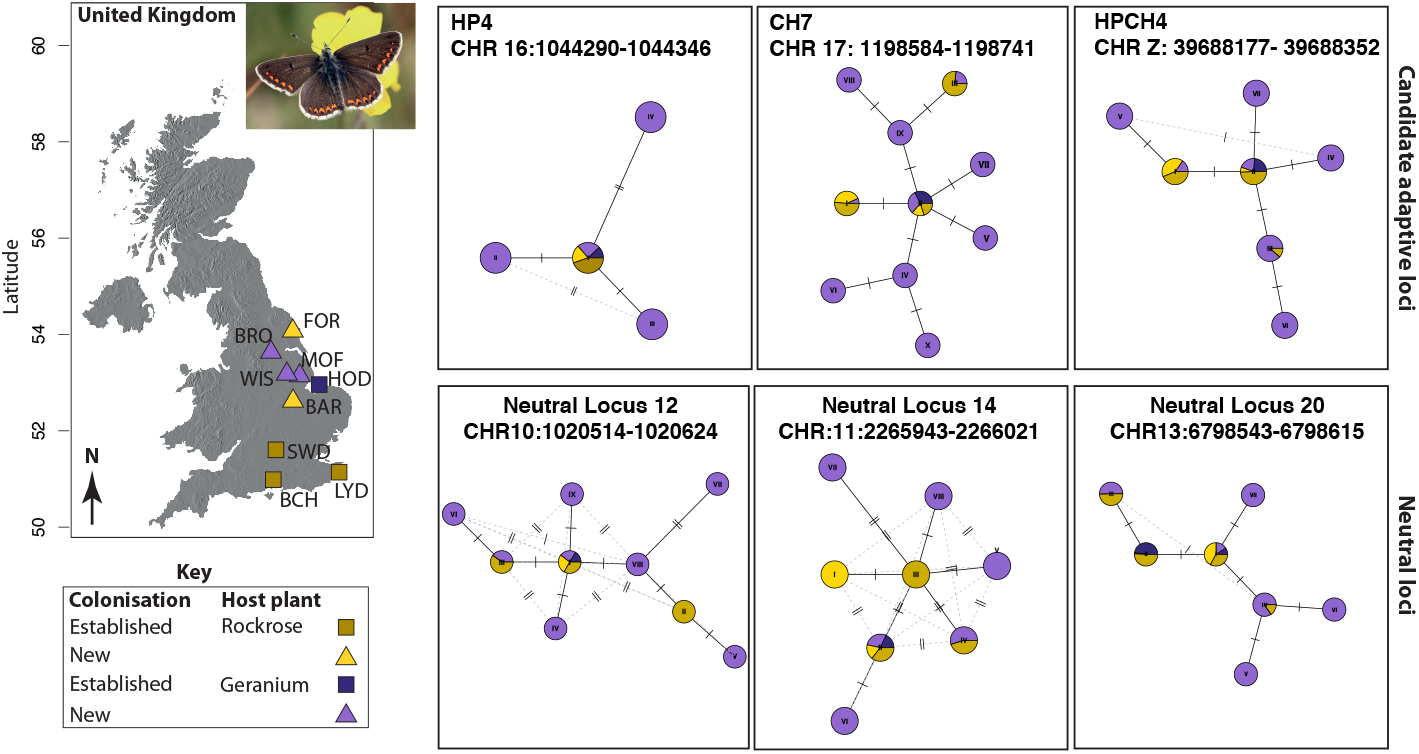
Haplotype network. Haplotype networks of a representative subset of the Host Plant (HP) outlier, Colonisation History (CH) outlier and Neutral loci. The remaining loci showed the same overall pattern and are shown in Figure S2. A genomic location of each haplotype is shown in Table S2, and haplotype frequencies in Table S3. Light yellow and light purple represent the newly established populations, while dark yellow represents the established South and dark purple represents HOD, the potential source of *Geraniaceae* adaptive haplotypes. The star-like configuration of haplotypes is indicative of a recent expansion, where haplotypes found exclusively in the new sites radiate from more common haplotypes found in the established range. If the adaptive haplotypes were introduced from HOD, we would expect the light purple haplotypes to radiate from dark purple haplotypes. Instead, the adaptive haplotypes originate either from the established South, or from both the established South and from HOD.

#### Reconstructing the colonisation history using a coalescent approach

The best-supported colonisation history model (Model3a) specified that New sites were colonised by an admixture of the established North-Eastern population (HOD) and the established populations in the South (Table 3; Fig 2). Point estimates of the model parameters suggests a slightly higher proportion of ancestry from the South (56%) than from HOD (44%) (Table 4), although this should be interpreted with caution based on the wide confidence intervals surrounding our estimates.

**Table 3.** Comparison of demographic models. Three demographic models (Figure 2) considering a) asymmetric migration between populations, and b) no migration after population divergence were simulated using FastSimCoal2. We report the log likelihood (log_10_L), number of parameters estimated (k), Akaike Information Criterion (AIC), rescaled AIC (ΔAIC), and weighting (wi) for each model. The best-supported model is highlighted in bold.

| Model | Log<sub>10</sub>L | k | AIC | deltaAIC | w<sub>i</sub> |
| --- | --- | --- | --- | --- | --- |
| 1a | -19917.8 | 15 | 91752.86 | 45.41 | 0.00 |
| 1b | -19944.37 | 9 | 91863.20 | 155.75 | 0.00 |
| 2a | -20007.05 | 15 | 92163.88 | 456.43 | 0.00 |
| 2b | -20019.58 | 9 | 92209.59 | 502.14 | 0.00 |
| **3a** | **-19907.07** | **17** | **91707.45** | **0.00** | **1.00** |
| 3b | -19933.18 | 11 | 91815.67 | 108.22 | 0.00 |

**Table 4.** Demographic parameters estimated from the best-supported coalescent model: Model3a. Point estimates were obtained from the run with the highest composite maximum likelihood. Upper and lower bounds were estimated from 100 bootstrap replicates of the model using these point estimates to construct the simulated site frequency spectrum. The parameter names are defined Figure 2 (model 3a), and priors are shown in the online supplementary material (Table S4).

| Parameter | Estimate | Lower Bound | Upper Bound | Category |
| --- | --- | --- | --- | --- |
| ANCSIZE | 544351 | 30800 | 1119249 | Population size |
| NNEW | 15562 | 5560 | 44915 | Population size |
| NSOUTH | 86206* |  |  | Population size |
| NHOD | 16659 | 8691 | 65541 | Population size |
| TADMIX | 9511 | 3680 | 13034 | Time |
| TPLUSDIV | 18372 | 4564 | 19874 | Time (complex parameter) |
| MIG01 | 2.31E-04 | 5.71E-10 | 6.56E-04 | Migration |
| MIG10 | 1.57E-08 | 3.82E-12 | 7.31E-05 | Migration |
| MIG02 | 4.78E-07 | 3.67E-11 | 3.30E-04 | Migration |
| MIG20 | 2.35E-05 | 3.51E-10 | 4.44E-04 | Migration |
| MIG12 | 1.45E-04 | 3.30E-11 | 7.67E-05 | Migration |
| MIG21 | 8.97E-05 | 1.19E-11 | 3.22E-04 | Migration |
| MUTRATE | 3.53E-09 | 3.26E-09 | 5.47E-09 | Mutation rate |
| RESIZE1 | 5.54 | 1.92 | 15.50 | Relative change in population size |
| RESIZE2 | 1.07 | 0.56 | 6.27 | Relative change in population size |
| MIGTOM | 0.56 | 0.11 | 0.95 | Migration |
| RESIZE3 | 6.31 | 0.36 | 12.98 | Relative change in population size |
| TDIV1 | 27883 | 11227 | 29246 | Time |

\*NSOUTH population size was fixed, and all other population sizes were estimated relative to NSOUTH.

## Discussion

### Rapid adaptation occurs by formation of genotypes from alleles across the range

The population genomic analyses presented here suggest that the rapid evolution of host plant use at the expanding range edge in *A. agestis* is associated with selection on genomic variation found across the species’ range and facilitated by high levels of gene flow between populations. Our analyses of haplotype data (Figure 5) and fastSimCoal2 simulations best support a scenario where climate adaptation occurred through evolution *in situ* during range expansion, rather than through colonisation by pre-adapted genotypes from coastal populations that already specialise on the new host plant. This finding is supported by the fact we find little evidence for a reduction in genome-wide genetic variance at newly-colonised sites, despite evidence that female oviposition preference has narrowed in the new habitats (Buckley & Bridle 2014). Buckley and Bridle (2014) also provide evidence for reduced fitness of long-established Geraniaceae-using populations on the Norfolk coast, when transplanted to *Geranium* sites in Lincolnshire, suggesting that established forms of Geraniaceae use may involve different traits to those associated with climate-driven range expansion.

FineRADstructure plots, and the lack of structure in the haplotype networks both in the adaptive and neutral loci, suggest ongoing gene flow from source habitats, or subsequent gene flow following an initial bottleneck associated with colonisation. Range expansion in *A. agestis* seems to occur by infilling of new habitats rather than a mass northward expansion *per se*, and with historical and ongoing high connectivity between neighbouring (source) populations. The lack of a strong signal in population size associated with range expansion could also be explained by a polygenic genomic architecture associated with the host-plant shift, meaning that selective sweeps during adaptation to novel conditions have had little effect on effective population size (see below).

### Shifts in host plant use associated with climate-driven range expansion

Although *A. agestis* females from the core range typically prefer Rockrose in field host choice experiments (Bridle et al 2014), they can also oviposit on Geraniaceae. However, the expansion into northern England has been associated with the loss of Rockrose use, certainly for Lincolnshire populations (Buckley and Bridle, 2014). Presumably therefore, some cost to maintaining preference for both host plants is responsible for the loss of rockrose laying preference in habitats where only Geraniaceae is present. Geographic variation in host plant choice has been found in several polyphagous butterfly species (e.g. Hanski *et al*. 2002; Nygren *et al*. 2006; Stålhandske *et al*. 2016), and these are driven by trade-offs such as differences in the host plant defence chemicals, host plant morphology and life history (e.g. phenology), or the regional and local abundance of host plants. Importantly, southern populations of Brown Argus continue to use both plant species, especially in warm years when *Geranium* becomes common at the margins of rockrose habitat. This may maintain the ability of females to use both species of host plant, especially if *Geranium* host plants provide more productive larval hosts than rockrose, at least in years when summers are relatively warm and dry (Stewart et al. 2021, and in press). By contrast, use of rockrose at these sites may be maintained by cold and wet years, when eggs laid on these plants should show substantially lower mortality than those laid on *Geranium* plants (<u>Stewart et al, in press</u>). Rockrose dominated sites predominate at the existing northern range margin, which could be a barrier to further expansion given the loss of rockrose use at new sites documented by Buckley and Bridle (2014). However, two sites included here constitute new rockrose sites; Forden is the northernmost site at the extreme northern range margin, and Barnack is the southermost newly colonised site. There is some evidence for rapid local adaptation (10-12 generations) from *Geranium* to rockrose preference at Barnack (Bridle et al. 2014; Buckley and Bridle 2014), driven either by evolution *in situ*, or re-colonisation from rockrose favouring individuals from the range core. The latter is more likely, given high levels of gene-flow found across the entire range (Fst=0.031), and the genetic similarity of new rockrose sites despite the geographic distance between them (Figure 1, Figure 3). An alternative explanation for the rockrose dominant site at the northern range edge (FOR) is that it is an established *A. agestis* site which hadn’t previously been detected. However, our assumption that Forden is a newly-established *A. agestis* site is supported by our genetic evidence: FOR clusters with the newly colonised sites in the fastStructure and fineRADstructure analyses and the PCA.

### The genomic basis of adaptation associated with range expansion

Using ddRAD markers allows us to investigate the genetic basis of adaptation associated with climate-driven range expansion at many loci, an important extension to previous AFLP data (Buckley et al 2012). In the first instance, it is remarkable that so few loci (and most with relatively low Fst levels for the outliers) are associated with the host plant shift, given that the plants are from different families. It could be that the shift in host plant is not as phenotypically demanding as we imagine, with potentially substantial overlap in the butterfly traits needed to sustain both interactions. Instead, the difference in microclimate between host-plants could be the main factor driving evolutionary change, demanding adaptation to cope with different pathogen abundance associated with each plant, or shifts in egg production or composition to alter desiccation or thermal resistance (see e.g. Stewart et al. 2021).

Alternatively, genomic variation already present and affecting host plant use in the established populations may mean that only small changes in allele frequency across many loci are needed to specialize on Geranium plants during expansion (see e.g. Pritchard *et al*. 2010). *A. agestis* is known to utilise *Geraniaceae* in the established part of the range where it is available, although they prefer *H. nummularium* in choice experiments. In addition, laboratory rearing of eggs (in warm conditions) is more successful on Geranium host plants than on rockrose, which seem to present a less nutritious (albeit more climatically reliable) host for larvae (Stewart et al. in press). We note also that we know nothing currently about local adaptation in the northern populations of these host plants, or any evolutionary responses that may have occurred in response to the novel invasion of Brown Argus into their communities. It seems likely that alleles determining use of a given host plant in Brown Argus will vary across its geographical range, possibly in response to plant local adaptation, and in relation to local microclimate (Stewart et al, 2021), as has been suggested in the Glanville Fritillary in Aaland (de Jong et al. 2018).

Two of the loci associated with host plant choice were located on the X-chromosome (Figure 4), which corresponds with evidence that the X-chromosome is important in Lepidopteran speciation driven by host-plant differences (Sperling 1994; Prowell 1998; Janz 2019). Within species variation in female oviposition choice has also associated with loci on the X-chromosome (e.g. in the comma buttefly, Nygren *et al*. 2006). Theoretically, genes on the X-chromosome could evolve faster than genes on autosomes, because that the recessive genes will be more available for selection to act on in the heterogametic sex (Charlesworth *et al*. 1987). It follows that a rapid change in female oviposition preference would be most successful if the genes were located on the X-chromosome. In contrast, a multi-species comparison of genes associated with host-plant preference in Lepidoptera found that loci were distributed throughout the genome, with a core set of genes located on the autosomes found across all butterfly/plant pairs (Nallu *et al*. 2018). A BLAST search of our loci found no overlap between our outlier loci and these candidate genes.

The four loci associated with both colonisation history and host-plant prevalence are particularly interesting because individuals colonising new sites during range expansions are often high dispersers. Increased individual movement (and extended searching) is also associated with *Geranium* host plant use (Bodsworth 2002). In addition, Bridle et al (2014) provide evidence for the evolution of increased dispersal ability in *A. agestis* in newly-colonised sites in the UK. However, a BLAST search of the candidate regions associated with dominant host plant or colonisation history did not reveal anything of particular interest (Table S1).

### Understanding the evolution of biotic interactions under climate change

This study provides further evidence that (a) evolutionary responses have been necessary for climatic shifts in the Brown Argus butterfly in the UK, and that (b) such evolution has occurred *in situ*, and has involved shifts in polygenic traits, involving the creation and establishment of novel genotypes during the range expansion, rather than the colonisation of pre-existing genotypes from elsewhere in the range. The Brown Argus example is highly instructive, given that rapid evolutionary responses are likely to be necessary for climate adaptation in many species and communities characterised by specialist biotic interactions, and look likely to depend on the maintenance of gene flow across a species’ geographical range (Hoffmann *et al*. 2021). Increased environmental unpredictability as well as variation are key in shaping the life history and behavioural strategies that evolve during adaptation to novel climates (Hoffmann & Bridle 2021). In this context, the Brown Argus example is also important, given its rapid range expansion is likely to have reduced its capacity to cope with increasingly unpredictable conditions in coming decades, by favouring specialisation on a host plant that is only productive during clement years (Stewart et al, in press). We may therefore expect that, due to evolutionary responses associated with range expansion, UK Brown Argus populations in coming decades will be characterised by greater fluctuations in population size, and loss of genetic variation in coming decades (O’Brien et al. in press), at least until selection favours the re-establishment of rockrose preference at the edge of its range.

## Acknowledgements

MdJ was funded by a Marie Curie Postdoctoral Fellowship. AJvR was funded by a Swiss National Science Foundation Early Postdoc Mobility Fellowship (P2ZHP2_178363). The generation of genomic data was funded by a grant to JB and MdJ from the Biomolecular Analysis Facility (NBAF) of the UK’s Natural Environment Research Council (NERC). We thank Roger Butlin and Chris Thomas for useful discussions and advice on sampling and data analysis, and to Natural England, the National Trust, individual landowners, and XXXXXXX for permission to collect, and for assistance in the field.

## Data accessibility

Sequence data will be submitted to the SRA on NCBI. Data and scripts used for each analysis will be uploaded to Dryad.

## Author contributions

- MdJ devised the study, secured ERC and NBAF funding, conducted the sampling and molecular lab work for the population genomics dataset, and data interpretation, and helped write the manuscript.
- AJvR designed and conducted the bioinformatic and genomic analysis_and interpretation, secured NBAF funding, and helped write the manuscript.
- SW designed and conducted bioinformatic analysis and created the draft genome assembly.
- CJY conducted the molecular lab work for the draft genome assembly
- JRB devised the study, hosted MdJ and assisted with study and sampling design, fieldwork, data analysis and interpretation, and helped write the manuscript (with contributions from all authors)
- MB assisted with genomic analysis and data interpretation
- CJ co-hosted MdJ and assisted with sampling design, genomic analysis and data interpretation.

