## Supplementary material for "The evolution of novel biotic interactions at ecological margins in response to climate change involves alleles from across the geographical range of the UK Brown Argus butterfly": Table S1-5; Figure S1-4

**De Jong, van Rensburg *et al.* Supporting information for online publication**

***Supplementary methods and results***

*Reconstructing the colonisation history using a coalescent approach*

Adaptation to *Geraniaceae* at the new sites could have been facilitated either by the arrival of pre-adapted genotypes. The established north-eastern population (HOD) is predominated by *Geraniaceae* and its geographic proximity to the new sites makes it a likely source of such pre-adapted genotypes. To determine the most likely origin of colonists at the new sites we compared two demographic models with 1) the established southern populations (South) and 2) HOD as the source using the coalescent simulator fastSimCoal2 (Figure S4) (Excoffier *et al.* 2013). We removed all non-neutral loci identified by our tests for selection and we reduced linkage in the dataset by retaining only one SNP per 10kb. This reduced the dataset from 31,158 to 3,486 loci, similar to the number of loci retained when we retain only one SNP in 400bp (i.e. the ddRAD library insert size; 3,868 loci), which suggests that our ddRAD loci are located at least 10kb apart. Missingness can bias the estimated site frequency spectrum (SFS), thus to increase the number of loci in the final dataset we used ANGSD (Korneliussen *et al.* 2014). Briefly, ANGSD estimates the likelihood of the allele frequency at each site. We allowed for a minimum depth of 2X per individual per locus. ANGSD then estimates the most likely single (HOD) and multidimensional (South and New) folded SFS for each region based on the loci that occur in both datasets. Our final SFS were based on $<5000$ loci for each joint folded SFS. The small number of loci used in constructing the SFS is likely to result in a poor fit of the simulated SFS to the data, and hence similar likelihood measures between models. Thus we further attempted to resample data within populations to reduce missingness using easySFS.py (https://github.com/isaacovercast/easySFS). This approach resamples loci that have a user specified minimum number of genotypes across all samples. In this way missingness within an individual is circumvented and a full genotype matrix can be created. However, we were unable to increase the number of loci above 3000 using this approach. We attempted to construct the coalescent model with this dataset.

The effective population size was fixed for South in the model so that all parameters could be estimated relative to this value. The effective population size ($N_e$) was calculated from the mutation rate (μ) and nucleotide diversity (π). Nucleotide diversity was calculated across all variable and invariable sites in windows of 1kb using vcftools (--window-pi function; π=0.001). We assume a mutation rate of 2.9 x $10^{-9}$ per base per haploid genome per generation based on the only direct estimate of Lepidoptera mutation rates (*H. melpomone*, Keightley *et al.* 2014). The effective population size was calculated as $N_e = (\pi/4\mu)$; South=86206.89, and HOD=86206. For parameter estimation we assume a generation time of 0.5 years as *A. agestis* in Britain are bivoltine. Because of the uncertainty associated with these parameters, and the small SFS compared with the number of parameters estimated (15), we expect wide and overlapping confidence intervals and uncertainty in the parameter estimates. As such, this test was to determine the relative likelihood of the different demographic models, rather than to ascertain the absolute values of the estimated parameters. We ran 100 independent simulations of each model in fastSimCoal2. Each run comprised 100 000 coalescent simulations and 40 expectation maximisation cycles. All parameters and priors are documented in Table S4.

We evaluated the model fit based on the lowest log likelihood and Akaike's information criterion (AIC). Based on these criteria we were unable to distinguish between the two models (Table S5). This is likely attributable to the small number of loci used in constructing the joint SFS, which is a problem often encountered with large ddRAD-type datasets which have been sequenced across multiple lanes.

### Supplementary Tables and Figures

***Table S1 Pairwise $F_{ST}$***

Population pairwise $F_{ST}$ (Nei 1973). Subtext denotes population history and host plant: E/N = Established/New; C/G= Cistaceae/Geraniaceae. Population codes correspond to the sites in Table 1.

| | **BAR-*NC*** | **BCH-*EC*** | **BRO-*NG*** | **FOR-*NC*** | **HOD-*EG*** | **LYD-*EC*** | **MOF-*NG*** | **SWD-*EC*** | **WIS-*NG*** |
|---|---|---|---|---|---|---|---|---|---|
| **BAR-*NC*** | 0 | | | | | | | | |
| **BCH-*EC*** | 0.027 | 0 | | | | | | | |
| **BRO-*NG*** | 0.029 | 0.035 | 0 | | | | | | |
| **FOR-*NC*** | 0.030 | 0.033 | 0.033 | 0 | | | | | |
| **HOD-*EG*** | 0.026 | 0.026 | 0.034 | 0.034 | 0 | | | | |
| **LYD-*EC*** | 0.033 | 0.022 | 0.047 | 0.042 | 0.031 | 0 | | | |
| **MOF-*NG*** | 0.021 | 0.034 | 0.026 | 0.027 | 0.027 | 0.040 | 0 | | |
| **SWD-*EC*** | 0.025 | 0.016 | 0.033 | 0.030 | 0.024 | 0.022 | 0.032 | 0 | |
| **WIS-*NG*** | 0.018 | 0.034 | 0.019 | 0.022 | 0.027 | 0.035 | 0.013 | 0.032 | 0 |

***Table S2 Outlier and Neutral loci***

Characteristics of each of the outlier loci identified in for the Host Plant and Colonisation History tests, as well as a set of 16 neutral loci used as a comparison in the haplotype network. For each locus we show the contig and base pair position on the draft *A.agestis* genome, as well as the *H. melpomone* linkage group (H.mel LG) and position that the sequence mapped to. To construct the haplotype networks we extracted the entire contig sequence that contained one or more SNPs (nr SNPs). Networks were constructed from sequences with more than one SNP.

" = Information is the same as in the prior cell.

NA=Not Applicable

- = No information available because loci didn't map to H.melpomone.

1* = Loci with only one SNP were excluded from haplotype networks.

| Outlier name | Association | A.agestis contig | Sequence Length (bp) | A.agestis position | H.mel LG | H.mel position | nr SNPs |
|---|---|---|---|---|---|---|---|
| Locus 1 | Host Plant | scaffold 962 | 20670 | 19860 | 01 | 3418280 | 5 |
| Locus 2 | Host Plant | contig 11951 | 11219 | 10614 | - | - | 7 |
| Locus 3 | Both | contig 5345 | 16650 | 510 | - | - | 11 |
| Locus 3 | Host Plant | " | " | 511 | - | - | " |
| Locus 3 | Both | " | " | 587 | - | - | " |
| Locus 4 | Host Plant | contig 17378 | 8898 | 3882 | - | - | 1* |
| Locus 5 | Host Plant | contig 5407 | 16568 | 5785 | - | - | 11 |
| Locus 6 | Host Plant | contig 19343 | 8252 | 4711 | - | - | 6 |
| Locus 7 | Both | contig 3838 | 19036 | 3376 | - | - | 38 |
| Locus 7 | Host Plant | " | " | 3378 | - | - | " |
| Locus 7 | Both | " | " | 3382 | - | - | " |
| Locus 7 | Host Plant | " | " | 3408 | - | - | " |
| Locus 8 | Col Hist | contig_4712 | 17502 | 1181 | - | - | NA |
| Locus 9 | Col Hist | contig_16912 | 9060 | 5538 | - | - | NA |
| Locus 10 | Col Hist | contig_3360 | 20031 | 2654 | - | - | NA |
| Locus 10 | Col Hist | " | " | 2666 | - | - | NA |
| Locus 11 | Col Hist | contig_59667 | 2029 | 912 | - | - | NA |
| Locus 12 | Col Hist | contig_18281 | 8583 | 3083 | - | - | NA |
| Locus 13 | Col Hist | contig_1883 | 24468 | 23537 | - | - | NA |
| Locus 14 | Col Hist | contig_19564 | 8188 | 5748 | - | - | NA |
| Locus 14 | Col Hist | " | " | 5815 | - | - | NA |
| NeutralLocus1 | Neutral | contig_12160 | 11104 | NA | - | - | 8 |
| NeutralLocus2 | Neutral | contig_10272 | 12222 | NA | 18 | NA | 3 |
| NeutralLocus3 | Neutral | contig_18932 | 8375 | NA | 14 | NA | 3 |
| NeutralLocus4 | Neutral | contig_4506 | 17827 | NA | - | - | 7 |
| NeutralLocus5 | Neutral | contig_2229 | 23122 | NA | - | - | 6 |
| NeutralLocus6 | Neutral | contig_37637 | 4427 | NA | - | - | 1* |
| NeutralLocus7 | Neutral | contig_25996 | 6518 | NA | - | - | 3 |
| NeutralLocus8 | Neutral | contig_662 | 33788 | NA | - | - | 2 |
| NeutralLocus9 | Neutral | contig_38039 | 4372 | NA | - | - | 4 |
| NeutralLocus10 | Neutral | contig_75445 | 1001 | NA | - | - | 2 |
| NeutralLocus11 | Neutral | contig_1985 | 24031 | NA | - | - | 9 |
| NeutralLocus12 | Neutral | contig_30084 | 5671 | NA | - | - | 1* |
| NeutralLocus13 | Neutral | contig_31360 | 5447 | NA | - | - | 5 |
| NeutralLocus14 | Neutral | contig_3461 | 19856 | NA | - | - | 7 |
| NeutralLocus15 | Neutral | contig_916 | 30752 | NA | - | - | 1* |
| NeutralLocus16 | Neutral | contig_2334 | 22774 | NA | - | - | 1* |

***Table S3 Haplotype frequencies***

Haplotype frequencies for each of the Host Plant outlier and Neutral loci. The contig/scaffold location of each locus on the *A.agestis* draft genome (*A.agestis* loc) is shown, as well as the number of SNPs on the extracted sequence (nr SNPs). The haplotype name (Haplotype) and frequency in each of the four population categories is shown.

HOD - EG = HOD, Established Geranium
NG = New Geranium
NC = New Cistaceae
South -EC = Established Cistaceae

(see attached Excel spreadsheet)

***Table S4 FastSimCoal parameters and their priors***

We used 13 regular and two complex parameters (#name) in our fastSimCoal models. Prior distributions were uniform (#dist; unif) in all cases. The effective population size of the ancestral (ANCSIZE) and extant populations (NNEW, NSOUTH, NHOD) were all drawn from broad priors. The time to the most recent common ancestor (TDIV1) is known to have occurred in the recent past, thus we chose a narrow prior (10-100 years). The later population division (TDIV2) is known to have occurred some time before TDIV1, thus we specify a broad complex prior (TPLUSDIV) to occur before TDIV1. Migration between each population pair was modelled seperately (MIGxx). The mutation rate (MUTRATE) is based on known estimates for Lepidoptera.

```
//        Priors and rules file
//        *********************
[PARAMETERS]
//#isInt?  #name        #dist   #min        #max
//all N are in number of haploid individuals
        1  ANCSIZE      unif        1000    1000000  output
        1  NNEW         unif        1000    1000000  output
        1  NSOUTH       unif        1000    1000000  output
        1  NHOD         unif        1000    1000000  output
        1  TDIV1        unif          10        100  output
        1  TPLUSDIV     unif          10      10000  output
        1  MIG01        unif           0        0.5  output
        1  MIG10        unif           0        0.5  output
        1  MIG02        unif           0        0.5  output
        1  MIG20        unif           0        0.5  output
        1  MIG12        unif           0        0.5  output
        1  MIG21        unif           0        0.5  output
        0  MUTRATE      unif    2.90E-09   5.50E-09  output

[RULES]

[COMPLEX  PARAMETERS]
        0  RESIZE       =       ANCSIZE/NSOUTH  output
        0  TDIV2        =       TDIV1+TPLUSDIV  output
```

***Table S5 FastSimCoal results***

A comparison of the parameter estimates and Akaike Information Criterion score (AIC) of the two models tested in fastSimCoal. Based on the AIC and log likelihood scores (loglhood) we could not distinguish between the two models.

| ***Model 1*** | | ***Model 2*** | |
|---|---|---|---|
| **AIC** | 21021.64 | **AIC** | 21069.54 |
| **loglhood** | -4558.71 | **loglhood** | -4569.11 |
| **Parameter** | | **Parameter** | |
| ANCSIZE | 1.43E+05 | ANCSIZE | 1.95E+05 |
| NNEW | 2.23E+04 | NNEW | 4.89E+03 |
| NSOUTH | 1.25E+05 | NSOUTH | 2.34E+04 |
| NHOD | 1.55E+04 | NHOD | 4.70E+03 |
| TDIV1 | 2.99E+02 | TDIV1 | 1.33E+02 |
| TPLUSDIV | 2.93E+02 | TPLUSDIV | 2.80E+01 |
| MIG01 | 0.00E+00 | MIG01 | 0.00E+00 |
| MIG10 | 0.00E+00 | MIG10 | 0.00E+00 |
| MIG02 | 0.00E+00 | MIG02 | 0.00E+00 |
| MIG20 | 0.00E+00 | MIG20 | 0.00E+00 |
| MIG12 | 0.00E+00 | MIG12 | 0.00E+00 |
| MIG21 | 0.00E+00 | MIG21 | 0.00E+00 |
| MUTRATE | 4.39E-09 | MUTRATE | 4.29E-09 |
| RESIZE | 1.15E+00 | RESIZE | 8.35E+00 |
| TDIV2 | 5.92E+02 | TDIV2 | 1.61E+02 |
| MaxEstLhood | -4537.903 | MaxEstLhood | -4568.141 |
| MaxObsLhood | -4506.546 | MaxObsLhood | -4506.546 |

***Figure S1 A 21-mer profile of the reference genome reads***

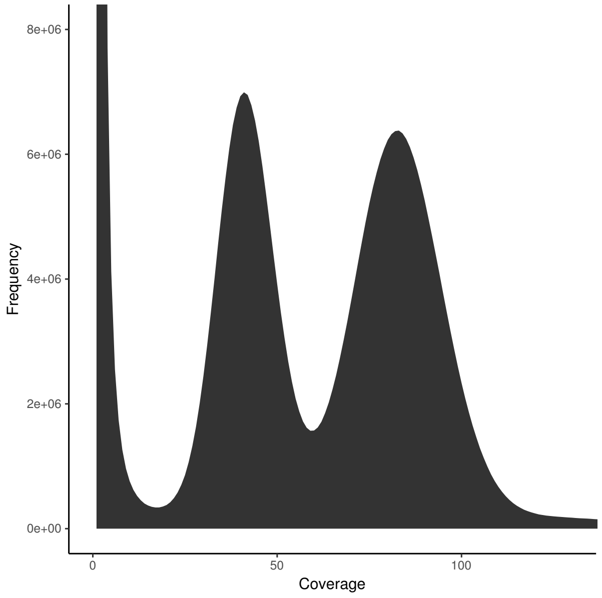


***Figure S2 Expected heterozygosity per population.***

We show the frequency distribution of expected heterozygosity per locus per population, as well as a box plot to show the median and distribution of the data.

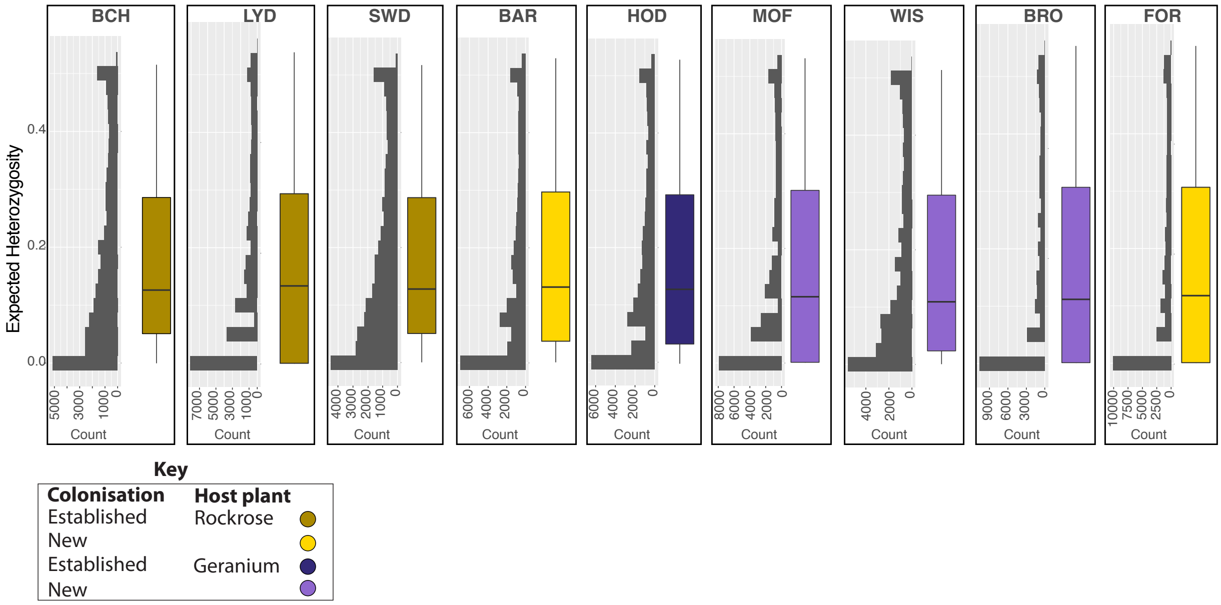

***Figure S3 Haplotype Networks***

Haplotype networks of the adaptive and neutral loci that are not included in Figure 2. Light yellow and light purple represent the newly established populations. If adaptive genotypes found in new *Geraniaceae* sites were introduced from the established *Geraniaceae* site (HOD, dark purple) we would expect light purple haplotypes to radiate out from a dark purple haplotype. Instead the haplotype networks of the adaptive loci closely resemble those of the neutral loci, with haplotypes largely co-occurring between established *Geraniaceae* and established *Cistaceae* sites, while haplotypes found exclusively in the new sites are derived from both established sites. Adaptive locus 7 is not displayed because of the large number of haplotypes (127), but results are summarised in Table S3. Contig numbers correspond to the draft *A. agestis* reference used in this manuscript.

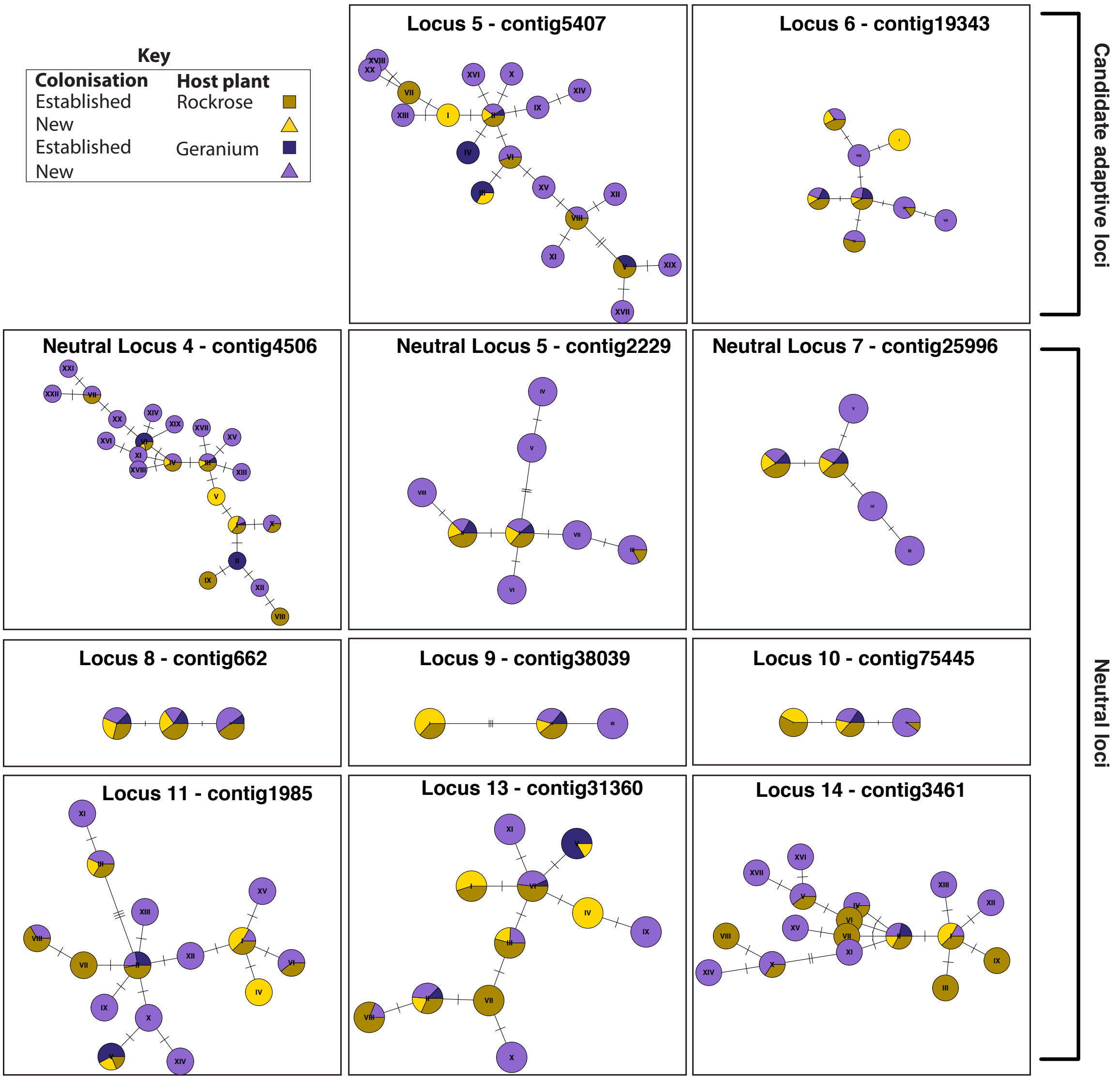

***Figure S4 Coalescent models to test colonisation history***

We tested for the most likely source population for the colonisation of the new sites using three coalescent scenarios in FastSimCoal2. We test if the New populations originated from the core southern populations (Model 1) or from the established population at Home Dunes specialised on *Geraniaceae* (Model 2). Effective population size (θ) was calculated for each population based on estimates of genetic diversity. Migration (m) and time to coalescence (TDIV), were drawn from uniform priors. All parameters and their priors are specified in Table S4.

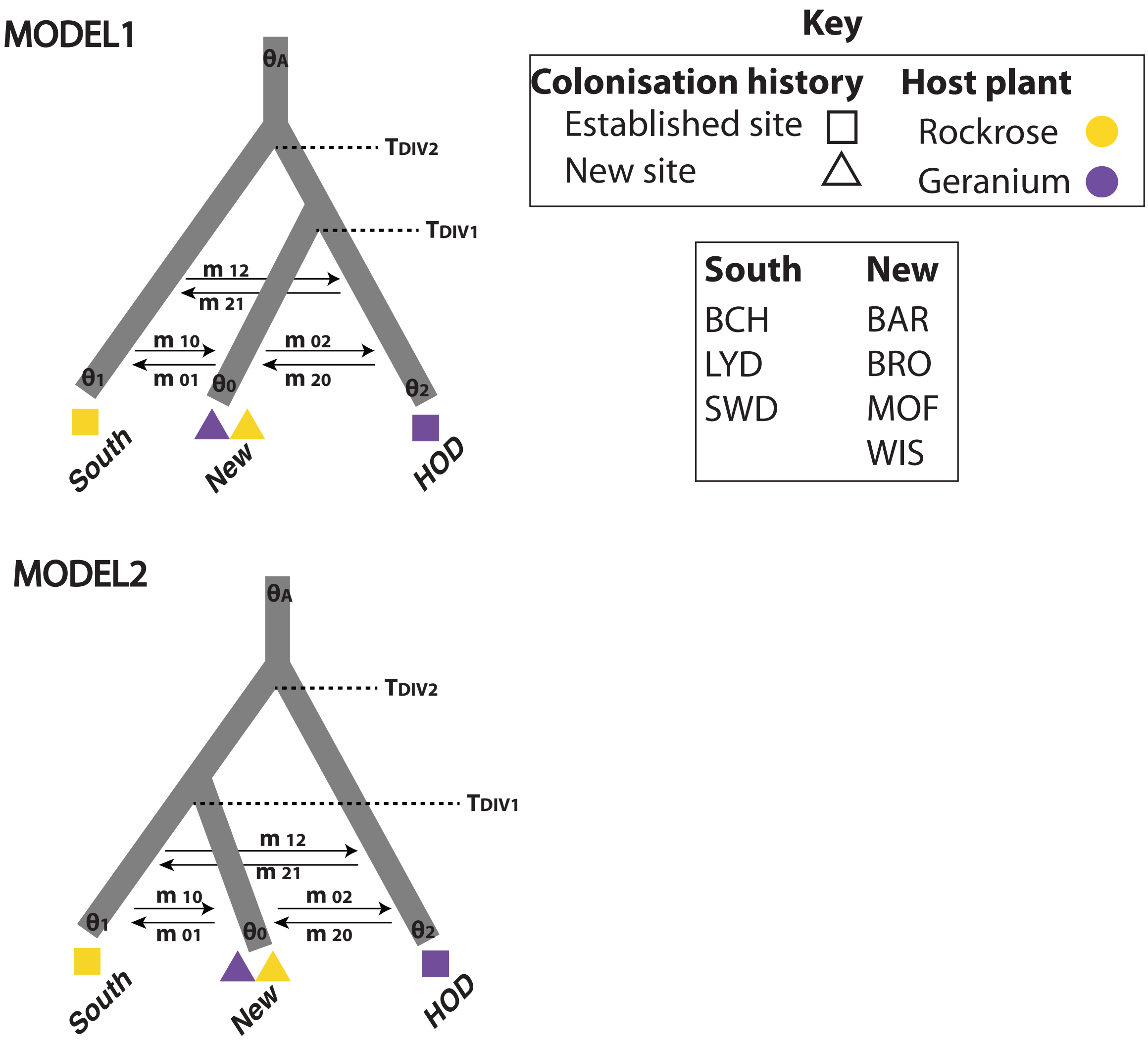
